# Postural analysis reveals persistent vigilance in paper wasps after conspecific challenge

**DOI:** 10.1101/2022.05.25.493496

**Authors:** Andrew W. Legan, Caleb C. Vogt, Michael J. Sheehan

## Abstract

Vigilant animals detect and respond to threats in the environment, often changing posture and movement patterns. In social animals vigilance is modulated not only by predators but also by threatening conspecifics. Precisely how social interactions alter vigilance behavior over time is not well understood. We report persistent effects of a simulated social challenge on the vigilance behavior of wild northern paper wasp foundresses, *Polistes fuscatus*. During the founding phase of the colony cycle conspecific wasps may usurp nests from the resident foundress, representing a severe threat. Using postural tracking, we found that after simulated intrusions wasps displayed increased vigilance during the minutes after the threat was removed. Sustained vigilance elicited after social threat manifested as increased movement, greater bilateral wing extension, and reduced antennal separation. However, no postural changes were observed after a control stimulus presentation. By rapidly adjusting individual vigilance behavior after fending off a conspecific intruder, paper wasp foundresses invest in surveillance of potential social threats, even when such threats are no longer immediately present. The prolonged state of vigilance observed here is relevant to plasticity of recognition processes as a result of conspecific threats.

## INTRODUCTION

Vigilance behavior in animals is demonstrated by changes in movement and body posture, famously in the still, bipedal stance of meerkat sentinels [1]. Movement and posture of specific body parts, especially the head and sensory organs, are primarily responsible for vigilance quality because they directly influence perception. For example, chaffinches turn their heads more after seeing a cat [2] and vigilant baboons blink less [3]. Animals must sometimes sacrifice vigilance quality in favor of other important activities, for example in the case of feeding juncos forfeiting some vigilance quality to lower their heads and eat [4].

Social animals, though characterized by their cooperative associations, face threats posed by conspecifics [5]. Recognition is an important mechanism mediating intraspecific aggression because encounters with different individuals and classes of individuals may impact fitness in distinct ways [6-10]. Social insects exhibit plasticity in nest guarding behavior in response to the frequency and valence of interactions with different classes of individuals (e.g., nestmates and non-nestmates) [11-14]. In response to encounters with non-nestmates, honeybees restrict admittance to the colony, sometimes rejecting their own nestmates [15,16]. These rejection errors are consistent with the signal detection theory concept of a shifting acceptance threshold [17]. With more frequent intruder encounters, the cost of accidentally accepting intruders increases, and the acceptance threshold is reduced to minimize acceptance errors. An alternative view considers variation in recognition behavior in terms of investment in recognition accuracy [18]. Recognition accuracy may be improved by persistent vigilant behavior of guards. Shifts in vigilance at the group level have been documented in honey bees, which allocate more guards at the colony entrance in response to threats [15,16,19]. How persistent vigilance is manifested in individual movement and posture has not been examined in a social insect.

To approach this question, we studied the northern paper wasp *Polistes fuscatus*. Paper wasps are ideal for field-based digital tracking because their unenveloped nest represents a fixed arena easily recordable by video. Automated tracking of wild foundress behavior is an as-yet unapplied tool for understanding the effects of intruder encounters on vigilance. During nest founding in the spring, *Polistes* foundresses guard the nest from conspecific wasps which may rob their brood or usurp their nests [20-25]. We simulated a guard context during the founding phase of single foundress *P. fuscatus* nests and leveraged digital tracking software to analyze wasp movement and posture.

## METHODS

We studied solitary *P. fuscatus* foundresses on their nests at the Liddell Field Station in Ithaca, NY (42°27’36.7” N, 76°26’39.2” W). In the spring of 2020, wild wasps initiated nests in modified wooden bird boxes (11.5 cm × 12.5 cm × 13.5 cm). Experiments were carried out from July 4^th^ to July 9^th^, 2020, before workers emerged, between 2PM and 9PM EST, during the active phase of wasps during peak summer. The mean nest size was 33 ± 8 (SD) cells. The experimental apparatus consisted of a 162.5 cm wooden dowel (7 mm diameter) guided through a 122 cm metal cylinder (1 cm diameter), taped to a step ladder (figure S1). The assays were video-recorded from below using a tripod-mounted Nikon D7200 camera with a Sigma Macro HSM lens and optical stabilizer (focal length: 105 mm; aperture: f/2.8).

Intruder wasps were collected from nests at a site (42°24’57.6” N, 76°31’22.6” W) 8.15 km southwest of the Liddell Station to ensure that foundresses had not encountered intruders before the experiment and were unlikely to be closely related [26]. Wasps were size matched to lures within 0.028 ± 0.013 grams (SD). Immediately before each simulated intruder trial, the intruder wasp was freeze-killed and fixed to a wooden dowel using an insect pin. Unique intruders were presented in each intruder trial. On a different day, each wasp was presented with the wooden dowel alone. All assays consisted of three 320 second intervals: pre-stimulus, stimulus, and post-stimulus. All nests were undisturbed, with experimental apparatus in place, for ≥ 5 min before beginning the pre-stimulus interval. During the stimulus presentation in both simulated intruder and wooden dowel trials, the stimulus was moved slightly by the experimenter at one-minute intervals to animate the stimulus. Three foundresses were excluded from analysis because a live intruder visited the nest during the experiment, and one foundress was excluded from analysis because it was accidentally flushed from the nest while setting up the experimental apparatus. Ultimately six foundresses were assayed.

We used SLEAP [27] to track seven points on the wasps: antennae tips, head, thorax-abdomen bridge (propodeum), abdomen tip, and wing tips (figure 1a). SLEAP was installed on a PC with a GeForce RTX 2080i graphics card. Videos were converted to gray scale and 20 frames per interval were manually labeled. We compared wing and antennae separation angle before and after stimulus presentations using paired t-tests. Statistical analysis was done using R version 3.6.1 [28].

**Fig 1.**
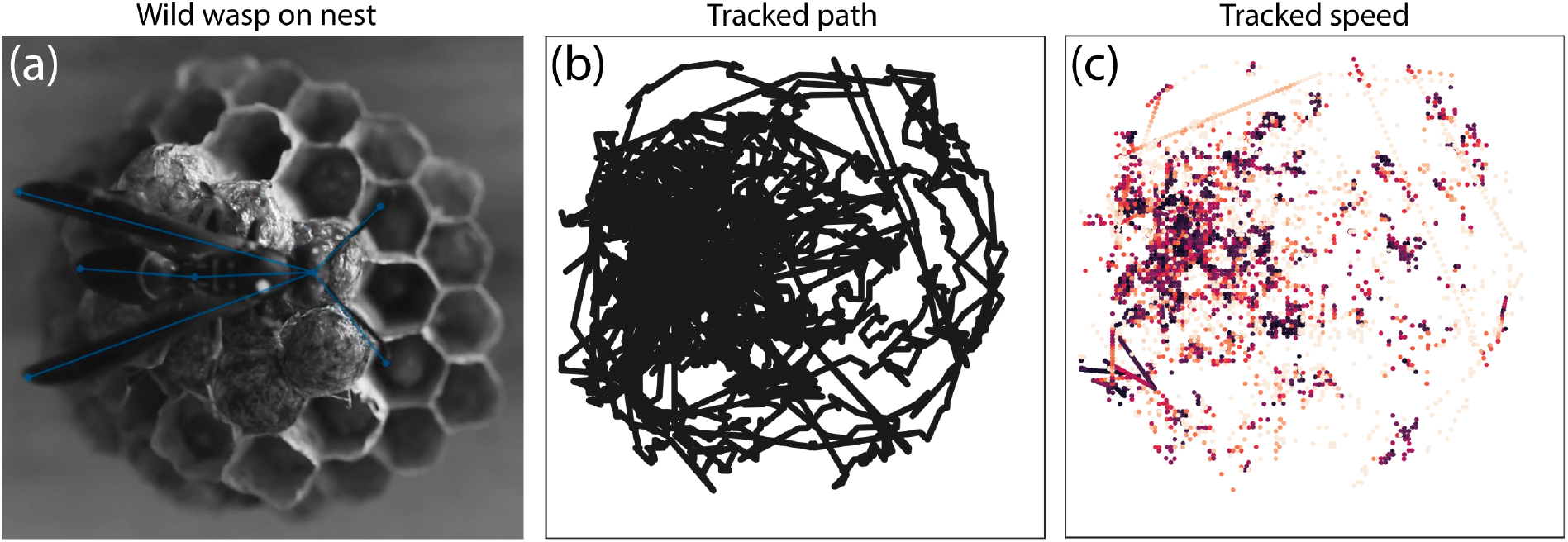
(a) A lone *Polistes fuscatus* foundress on the nest after a simulated intruder encounter. (b) Tracks of the position of the thorax of the wasp over a 320 second interval after simulated intrusion. (c) Points designate the position of the thorax and are color-coded by speed, with lighter color representing faster movement.

## RESULTS

When the lure was presented during simulated intruder trials, wasps responded by antennating the lure, then responded aggressively by biting, mounting, and stinging the lure (video S1). These are all stereotyped wasp aggressive behaviors [29-31]. In control stimulus trials, wasps investigated the dowel, including antennation and occasional mounting, but did not escalate aggression (video S1). SLEAP successfully tracked body parts in 84 ± 21% (SD) of frames across body parts before and after stimulus presentation (table S1).

Simulated intruder encounters caused persistent changes in movement and posture while control experiments did not. Exposure to the simulated intruder caused an increase in the total distance traveled after the intruder was removed (head: t = -2.5682, df = 5, P-value = 0.05014; thorax: t = -2.9614, df = 5, P-value = 0.03147; figure 2a). Dowel presentations did not lead to sustained increases in movement afterward (head: t = 0.27264, df = 5, P-value = 0.796; thorax: t = 0.037249, df = 5, P-value = 0.9717; figure 2a). Wing posture was affected by the simulated intruder. The mean wing extension angle after intruder encounter was significantly greater than before (t = -6.1917, df = 5, P-value = 0.001603; figure 2b). No significant change in wing extension angle was observed after the wooden dowel presentation (t = -1.2836, df = 5, P-value = 0.2556; figure 2b). There was a significant decrease in the mean antennal separation angle after intruder encounter (t = 4.1753, df = 5, P-value = 0.008695; figure 2c). No significant change in mean antennal separation angle was observed after the wooden dowel presentation (t = 0.40974, df = 5, P-value = 0.699; figure 2c).

**Fig 2.**
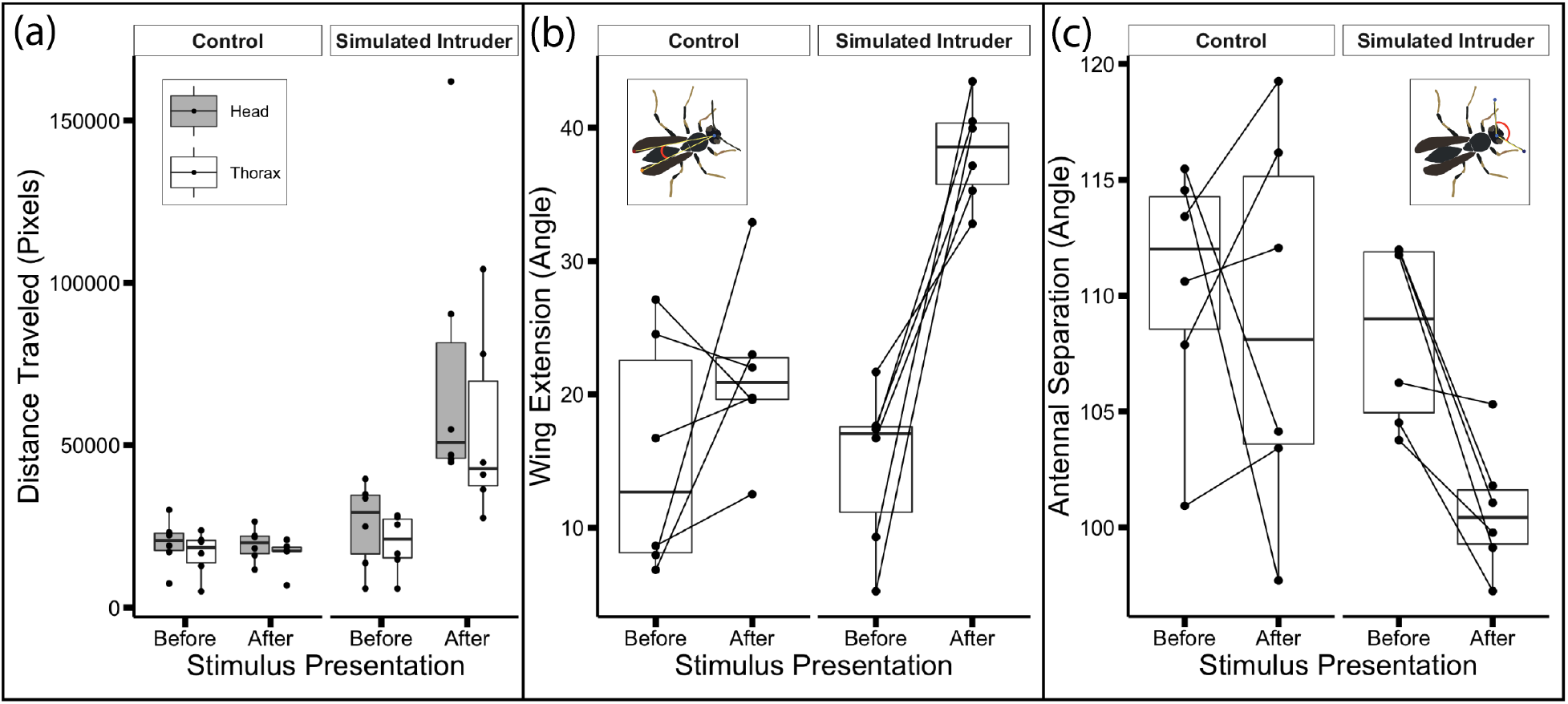
Box and whisker plots display comparisons of measures of movement and posture across trials. (a) Total distance traveled by head (gray) and thorax. (b) Wing extension angle. (c) Antennal separation angle.

The rapid and varied movement of wasps during simulated intruder presentations, and the presence of a second, pinned wasp, precluded successful digital tracking. However, digital tracking during the dowel presentations was feasible. During the dowel presentation, wasps did not move more than they did before the presentation, based on the total distance traveled by the thorax (t = -1.2475, df = 5, P-value = 0.2675; figure S2a). There was a significant increase in wing extension angle during the dowel presentation compared to before (t = -2.8063, df = 5, P-value = 0.03771; figure S2b). This increase in wing extension did not persist after the dowel was removed.

## DISCUSSION

Social challenge presented by simulated conspecific intruders elicited sustained vigilance in *P. fuscatus* spring foundresses. We leveraged computer vision to analyze wasp body posture in the field and found that sustained vigilance manifested in changes to wasp movement and posture.

Methods in automated tracking of behavior have been applied by scientists studying neurobiological mechanisms of animal movement and pose, collective behavior, and social interactions [32-35]. Automated tracking studies of insects are often carried out in controlled environments, which is feasible when the behavior of interest is robust to laboratory conditions. For example, digital tracking has been used to characterize the foraging behavior of hawkmoths *Manduca sexta* [36-38], and to characterize the wing kinematics of flies and honey bees as well as honey bee wing fanning behavior [39-41]. Complex social behaviors are less robust to experimental laboratory conditions, requiring field observations to draw reliable conclusions. But few studies have applied digital tracking of individual social insect posture in the wild (but see ref. 41).

*Polistes* paper wasps are ideal for computer vision-assisted digital tracking and pose estimation in the field. Compared to eusocial ants, bees, and hornets, *Polistes* societies remain relatively small, peaking at ∼135 cells [20]. *Polistes* colonies are generally single-layer nests. In terms of video recording, a drawback to this architecture is that there is usually space between the nest and the substrate to which it is fixed, so wasps can crawl out of view of the camera behind the nest. While the nest can be treated as two-dimensional for the purpose of digital tracking, the wasp’s body is not always parallel to this plane, leading to difficulties in tracking a wasp perched on the side of the nest. These challenges could be solved with multiple cameras recording the nest from different angles, as has been done recently for 3-dimensional tracking in laboratory rodents [42,43]. Another challenge for digital tracking is the rapid movement of wasps during the simulated intrusions, but cameras with faster frame rates could overcome this issue.

Natural threats that would induce nest-guarding behavior in solitary foundresses include intraspecific brood-robbing and nest usurpation [23-25]. In a study of multiple foundress *P. fuscatus* nests during the founding phase, natural encounters with intruders occurred about once per day, with intruders evicted within 40 seconds [25]. Three trials in our study were interrupted by natural intruders, highlighting the pervasive nature of conspecific threats for *P. fuscatus* foundresses. The 320 second lure presentation in our assays likely simulated a worst-case scenario for foundresses, akin to a nest usurpation attempt.

Postural changes displayed by vigilant wasps included wing extension and reduced antennal separation. Upon presentation with the simulated intruder, wasps approached the lure with outstretched antennae before reacting aggressively. In general, social insects utilize chemical cues to discriminate nestmates and non-nestmates [44-46]. While *P. fuscatus* wasps rely on vision to recognize individuals, nestmate recognition is mediated by olfaction [47,48]. The honeybee *Apis mellifera* responds to different odors with different antennal posture, depending on experience, demonstrating the function of antennal posture in perceiving odors [49,50]. Reduced antennal separation may indicate that wasps are orienting their antennae to detect chemical cues, such as the cuticular hydrocarbon signatures used by many social insects to discriminate nestmates and non-nestmates [51-54]. Visual cues may also be important in discriminating nestmates and non-nestmates at the early phases of the colony cycle, and the absence of nestmates may favor universal rejection [17,55].

Vigilant wasps moved more after fighting a simulated intruder, as measured by total distance traveled. This increased movement was observed throughout the 320 s interval after the simulated intruder was removed (figure S3). By moving throughout the nest surface, vigilant guard wasps may be better prepared to defend against an intruder approaching from any direction.

In *Polistes*, wing extension and antennal separation may be useful measures for studying how the social environment influences internal state. The reliable associations between unilateral wing-extension and courtship, and between bilateral wing extension and aggression, have been useful measures for studying the neural basis of aggression and courtship in the fly *Drosophila melanogaster*, especially the roles of P1 neurons in orchestrating persistent internal states causing aggression and courtship [56,57].

The internal state associated with vigilance in *P. fuscatus* may represent an emotional primitive, as defined by Anderson and Adolphs [58] as an internal state exhibiting scalability, valence, persistence, and generalization. Regarding scalability, we found evidence that wing extension can be ordered along a gradient corresponding to low vigilance (before stimulus), medium vigilance (during dowel presentation), and high vigilance (after simulated intruder presentation, demonstrating behavioral persistence) (figure S2). *P. fuscatus* vigilance behavior is associated with aggression towards conspecific intruders, suggesting negative valence. After social challenge, vigilance is persistent. More work is needed to assess the generalization of *P. fuscatus* vigilance behavior, for example by presenting wasps with neutral stimuli after social challenge.

Increased encounters with non-nestmate intruders can shift social insect recognition processes to become more exclusive, resulting in recognition error in the form of increased aggression towards nestmates [11,14-16,59]. From the perspective of signal detection theory, individual vigilance could be mechanistically related to acceptance threshold. If persistent vigilance and acceptance threshold shift are coupled, then there will be more aggression towards nestmates following intruder encounters. Alternatively, vigilance may affect recognition independent of acceptance threshold. For example, persistent vigilance may accompany increased investment in accurate recognition [18]. Evidence supporting this hypothesis may be found in the carpenter ant: exposure to alarm pheromone increased accuracy of both nestmate acceptance and non-nestmate rejection [60]. Persistent vigilance may therefore increase recognition accuracy, while the acceptance threshold is shifted depending on non-nestmate encounter rates [17]. Future work should explore how individual wasp vigilance relates to shifts in nestmate recognition processes.

## Supporting information

AppendixS1

tableS1csv

DSC_8565_1.avi.predictions.cleaned.analysis.h5_tracks

DSC_8565_3.avi.predictions.cleaned.analysis.h5_tracks

DSC_8569_1.avi.predictions.cleaned.analysis.h5_tracks

DSC_8569_2.avi.predictions.cleaned.analysis.h5_tracks

DSC_8569_3.avi.predictions.cleaned.analysis.h5_tracks

DSC_8571_1.avi.predictions.cleaned.analysis.h5_tracks

DSC_8571_2.avi.predictions.cleaned.analysis.h5_tracks

DSC_8571_3.avi.predictions.cleaned.analysis.h5_tracks

DSC_8572_1.avi.predictions.cleaned.analysis.h5_tracks

DSC_8572_3.avi.predictions.cleaned.analysis.h5_tracks

DSC_8573_1.avi.predictions.cleaned.analysis.h5_tracks

DSC_8573_3.avi.predictions.cleaned.analysis.h5_tracks

DSC_8574_1.avi.predictions.cleaned.analysis.h5_tracks

DSC_8574_2.avi.predictions.cleaned.analysis.h5_tracks

DSC_8574_3.avi.predictions.cleaned.analysis.h5_tracks

DSC_8601_1.avi.predictions.cleaned.analysis.h5_tracks

DSC_8601_2.avi.predictions.cleaned.analysis.h5_tracks

DSC_8601_3.avi.predictions.cleaned.analysis.h5_tracks

DSC_8604_1.avi.predictions.cleaned.analysis.h5_tracks

DSC_8604_3.avi.predictions.cleaned.analysis_tracked

DSC_8605_1.avi.predictions.cleaned.analysis.h5_tracks

DSC_8605_2.avi.predictions.cleaned.analysis.h5_tracks

DSC_8605_3.avi.predictions.cleaned.analysis.h5_tracks

DSC_8607_1.avi.predictions.cleaned.analysis.h5_tracks

DSC_8607_2.avi.predictions.cleaned.analysis.h5_tracks

DSC_8607_3.avi.predictions.cleaned.analysis.h5_tracks

DSC_8608_1.avi.predictions.cleaned.analysis.h5_tracks

DSC_8608_3.avi.predictions.cleaned.analysis.h5_tracks

DSC_8609_1.avi.predictions.cleaned.analysis.h5_tracks

DSC_8609_3.avi.predictions.cleaned.analysis.h5_tracks

README

## DATA ACCESSIBILITY

Videos are available on Zenodo: https://doi.org/10.5281/zenodo.6582229. The electronic supplementary material includes outputs of tracking (30 csv files), table S1 (1 excel file and 1 csv), video S1 (1 mp4 file) and Appendix S1 containing figures S1-S3 (1 pdf file).

## FUNDING

This work was supported by National Science Foundation [Graduate Research Fellowship Program DGE-1650441 to AWL, CAREER grant DEB-1750394 to MJS], Cornell University [Neurobiology and Behavior Departmental Grant to CCV], and National Institutes of Health [DP2-GM128202 to MJS], and by the North American Section of the International Union for the Study of Social Insects (IUSSI) Robert L. and Louise B. Jeanne Social Wasp Research Grant [to AWL].

## AUTHOR CONTRIBUTIONS

AWL conceptualized and carried out field experiments, carried out analyses, and wrote the paper. CCV manually labeled frames and used digital tracking software to generate tables of tracked body parts. All authors contributed to conceptualizing analyses. All authors helped edit the manuscript.

## Notes

https://doi.org/10.5281/zenodo.6582229

### Competing Interest Statement

The authors have declared no competing interest.

