## AppendixS1 for "Postural analysis reveals persistent vigilance in paper wasps after conspecific challenge"

**APPENDIX S1 FOR**

### Postural analysis reveals persistent vigilance in paper wasps after conspecific challenge

Andrew W. Legan[a], Caleb C. Vogt[a], Michael J. Sheehan[a]

a. Laboratory for Animal Social Evolution and Recognition, Department of Neurobiology and Behavior, Cornell University, Ithaca, NY

| **CONTENTS** | **PAGE** |
|---|---|
| Figure S1: Experimental apparatus | 1 |
| Figure S2: Box and whisker plots display measures of movement and posture | 2 |
| Figure S3: Heatmaps showing speed of tracked body parts over time | 3 |

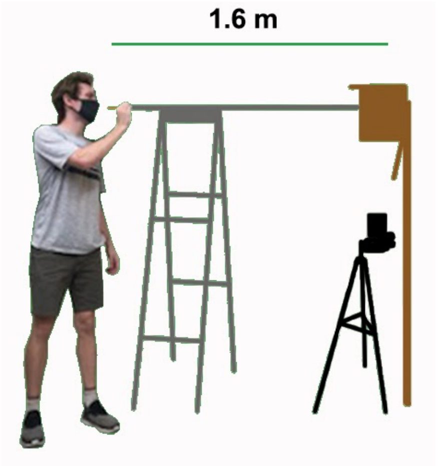


**Figure S1**. Experimental apparatus.

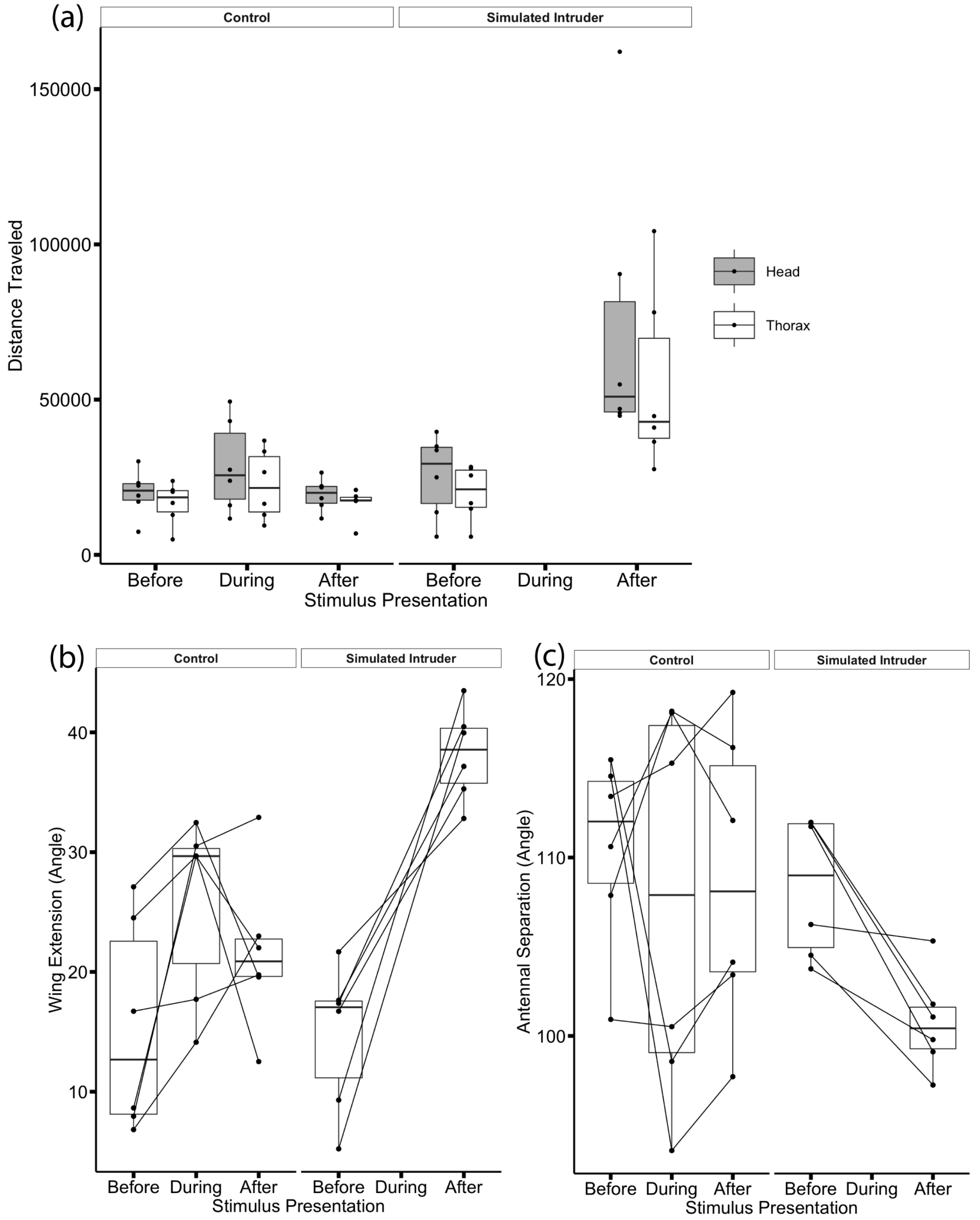


**Figure S2**: Box and whisker plots display comparisons of measures of movement and posture across trials. Same data as Figure 2 of main text, except interval 2 of control trials has been tracked. (a) Total distance traveled by head (gray) and thorax in each interval. (b) Wing extension measured as an angle in degrees. (c) Antennal separation measured as an angle in degrees. In each plot, the horizontal line marks the median of the data points, the box brackets the middle fifty percent of data points, and the whiskers extend to the smallest and largest values at most 1.5 * the interquartile range.

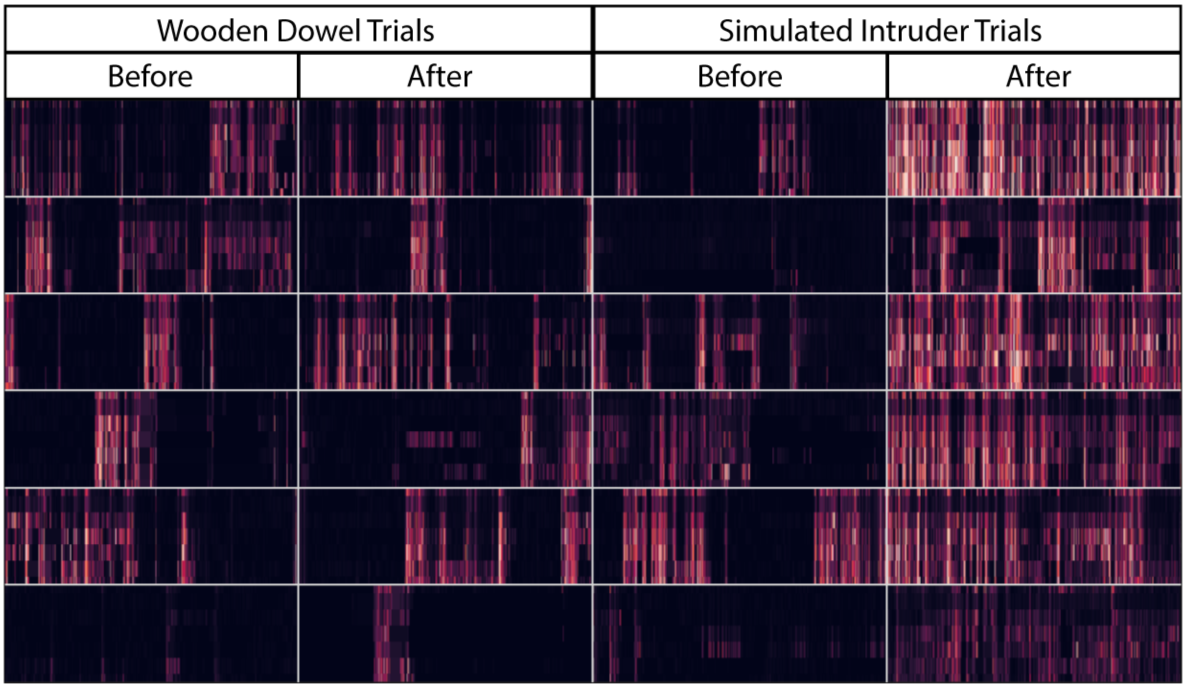


**Figure S3**: For six wasps assayed in two experimental intervals of twelve trials, the speeds of seven tracked body parts over time are shown in a heatmap. Six rows of heatmaps correspond to six individual wasps presented with a wooden dowel and an intruder wasp on separate days. Rows within each heatmap correspond, from top to bottom, to seven tracked body parts: head, thorax-abdomen bridge (propodeum), abdomen tip, left wing tip, right wing tip, left antenna tip, and right antenna tip.
