## Supplementary material for "Postural analysis reveals persistent vigilance in paper wasps after conspecific challenge": README

# README

This document describes the videos uploaded to bioRxiv under the title "Postural analysis reveals persistent vigilance in paper wasps after conspecific challenge."
Study was located in Ithaca, NY in July, 2020.
Please contact Andrew Wesley Legan at with questions.

## "AppendixS1.pdf"
1.1MB
Appendix file containing three supplementary figures.

## "tableS1.xlsx"
13KB
Table of video metadata, tracking quality, and movement/posture measurements.

## "tableS1.csv"
4KB
Same as above in comma separated format.

## "videoS1"
1080p 
89.2MB
Sequence shows 250 frame excerpts of videos of all wasps assayed before, during, and after dowel and simulated intruder presentations, respectively. 

## "DSC_...avi.predictions.cleaned.analysis.h5_tracks.csv"
72 csv files
64MB together
x and y coordinates of seven body parts tracked using SLEAP.
